# Context-dependent oviposition reveals strong association between acceptance and preference in the Mediterranean fruit fly

**DOI:** 10.1101/2025.06.20.660717

**Authors:** Benoît Facon, Madeline Chauve, Virginie Ravigné, Bruno Serrate, Myriam Robejean, Antoine Fraimout, Julien Foucaud

## Abstract

Oviposition behavior in phytophagous insects is influenced by different stimuli and plays a key role in pest dynamics and crop loss. This study used 3D-printed artificial fruits varying in colour (yellow, blue, white) and odour (cherry, orange, banana) to test how visual and olfactory cues affect oviposition acceptance (no-choice) and preference (choice). In no-choice assays, the nine artificial fruits displayed sufficiently different visual and olfactory cues to trigger different egg-laying outputs (by a factor 1:3 between the least attractive fruit, white fruit with banana scent and the most attractive fruit, yellow fruit with cherry scent). While cues acted independently in no-choice settings, significant interactions were observed in choice conditions, highlighting multimodal sensory integration. In choice assays, the number of eggs laid and female preference depended on both the characteristics of fruits and their context. However a strong correlation was found between acceptance and preference. The relationship found between acceptance and preference implied that when a fruit seemed preferred in no-choice assays, it was even more preferred in one-choice assays. We finally discussed the practical implications for behavior-based pest management strategies.

## Introduction

Phytophagous pest insects are of particular importance for food safety as they are responsible for between 20 and 40% of crop production losses worldwide (Culliney 2014, Deutsch et al. 2018). For a large number of these species, larvae are unable to move from one plant to another to find the best conditions for their successful growth and development. Therefore, the choice of the host plant in which larvae eventually develop relies on gravid females and their oviposition behavior. Consequently, phenotypic variation in females’ oviposition behavior is tightly linked to offspring growth and survival, and constitutes an important fitness component on which selection can act (Jaenike 1978, Cunningham 2012). Aside from its relevance in the field of evolutionary ecology (see for instance Futuyma & Moreno 1988, Bernays & Chapmann 1994), the study of insect oviposition behavior is particularly important for the management of pest species. In an effort to reduce the use of synthetic chemical pesticides, studying oviposition behavior is needed for the development of behavior-based pest control techniques such as masking of host plant compounds to disrupt recognition, selecting crop varieties emitting less attractive volatiles, using trap crops or combining repellent and attractant elements in push–pull systems (Anton & Cortesero 2022). Integrating evolutionary theory to studying oviposition behavior seems therefore to be a promising approach to determine the efficiency of alternative pest control techniques, and the evolutionary response of insect populations to this new form of selection (Sentis et al. 2022).

Oviposition behavior involves a complex suite of traits. Insects locate and discriminate among plants by a variety of sensory means that ultimately lead to plant acceptance or rejection (Jaenike 1978). Even though olfaction seems sufficient to locate and identify a suitable host for some phytophagous species, other senses, i.e., vision and contact chemoreception may also play roles in this process (Tasin et al. 2011, Bernays & Chapman 1994, Balkenius et al. 2009), with cross-stimuli interactions being the rule rather than the exception. The whole fruit display, involving scent, colour, pattern, shape, touch, and taste, is perceived by sensory systems and integrated in higher brain centers to trigger the correct behavioral response (Riffell 2020, Saitta et al. 2023).

Recent studies have highlighted non-trivial interactions between stimuli (mainly visual and olfactory). For instance, some insects such as the melon ladybird, *Chnootriba elaterii*, cannot discriminate plants based solely on visual or olfactory signals (Saitta et al. 2023). They need to perceive both cues simultaneously to select their host plant. When sensory cues come into conflict, i.e., visual stimuli do not match olfactory stimuli, then insect response can be altered in a highly variable manner depending on the odours and colours involved (Yv et al. 2015, Riffell 2020). Just as the suite of traits involved in oviposition behavior, the stimuli produced by the plants are fundamentally multivariate, with each stimulus potentially varying in its mode of action. Accounting for these multimodal interactions is therefore important to understand oviposition choice of phytophagous insects (Balkenius & Balkenius 2016). Finally, in other insects, the olfactory input modulates the visual system, whereas the visual system does not modulate the olfactory responses (Riffell 2020).

The response to stimuli is also highly context-dependent, depending in particular on the current availability of different host species (Schäpers et al. 2017). For instance, *Drosophila melanogaster* females prefer to oviposit on ethanol-containing substrates, but only as long as they are very close to an ethanol-free substrate (Sumethasorn & Turner 2016). The importance of the environmental context has led to the definition of two quantitative traits: oviposition acceptance and oviposition preference (Fanara et al. 2023). Acceptance is defined as the propensity of the female to lay eggs on a particular host when it is the only available possibility, as typically measured by no-choice experiments. In the field, this would correspond to monocultures where all plants produce the same stimuli. Preference is the propensity of the female to lay eggs on a host when other hosts are available. Preference is best assessed through one- or multiple-choice assays. In the field, such situations would be found in mixed cultures or in natural plots harboring a diversity of host plants. Resolving the relationship between oviposition acceptance and preference is particularly important for the development of evolutionary-informed pest control strategies. Several studies have tested this relationship using different insect species with contrasting results (Yang et al. 2015, Charlery de la Masselière et al. 2017, Olazcuaga et al. 2019, Fanna et al. 2023). Charlery de la Masselière et al. (2017) found that acceptance and preference showed a significant positive correlation for four Tephritidae species out of six studied species. On the contrary, Fanara et al. (2023) found that preference is not correlated to acceptance in *Drosophila melanogaster*, suggesting that these behaviors are independent.

To our knowledge, no study to date has simultaneously quantified acceptance and preference while experimentally disentangling the relative contributions of both visual and olfactory stimuli. By integrating these four components within a unified framework, our study provides a comprehensive assessment of how multimodal cues shape oviposition decisions and how acceptance and preference interact to determine final host-use outcomes using the Mediterranean fruit fly *Ceratitis capitata* as a study model. Belonging to the Tephritidae family (true fruit flies), *C. capitata* is a multivoltine generalist species with more than 300 described host plants (Dias et al. 2018). Originating from tropical sub-Saharan Africa, it now has a global distribution and is considered as one of the most important insect pests for fresh fruit and vegetables production worldwide because of both direct damage to fruits and quarantine regulations (USDA 2006). Females lay their eggs in ripening or ripe fruits, leading to accelerated maturation and premature fruit drop, rendering them unfit for consumption and unsuitable for marketing (Dias et al. 2018). To study oviposition choice in *C. capitata*, we produced 3D-printed oviposition supports to mimic fruits that can vary in colour and odour, two essential cues for host choice in *C. capitata*. In this study, we measured the behavior of female flies in response to three colours (blue, yellow, and white) and three odours (cherry, banana, and orange). We used a full-factorial design to thoroughly determine the impact of visual and olfactory cues on the oviposition behavior of *C. capitata* in two types of environments: no-choice and one-choice assays, allowing to respectively evaluate oviposition acceptance and preference. By delving into the intricacies of the oviposition behavior, we aim at providing valuable insight into the determinants of a major trait of insect pest biology and hope to contribute to guide efforts in agroecological crop protection.

## Materials and Methods

### Study system and experimental conditions

The individuals used in this study were derived from a population reared in the arthropod rearing and phenotyping facilities of the Biology Centre for Population Management (Montpellier, France) for two years following crosses between individuals collected in the wild between 2021 and 2022 from three different southern France localities: San-Giuliano (Corsica), Torreilles (Pyrénées-Orientales), and Millas (Pyrénées-Orientales). Both rearing and experiments were conducted in climate chambers under controlled conditions (temperature: 25 ± 1°C, humidity: 60 ± 10%, and photoperiod: 12L:12D). For the mass rearing of this population, white mesh cages of 30 × 30 × 30 cm containing approximately 1,800 individuals with a sex ratio of 1:1 were used. Larvae were reared on an artificial diet composed of dehydrated carrot powder, brewer’s yeast, dehydrated potato, water, and Nipagin/Sodium benzoate. In laboratory conditions, the developmental cycle from egg to adult emergence lasted approximately 17 days. These cages allowed the creation of cohorts of gravid females every week for the experiments. After adult emergence, females had the opportunity to mate for approximately five days before being isolated in experimental cages. The experiments were conducted in smaller white mesh cages of 15 × 15 × 15 cm (BugDorm-4M1515 insect cage that has nylon netting (44 × 32 mesh) with 700 µm aperture). Each cage, representing a replicate, contained eight reproductive females of the same age (five days).

Artificial fruits were chosen over real fruits for three main reasons. Firstly, fruit availability varies seasonally. Additionally, the inter-fruit variability within a species may be high. Lastly, previous studies have shown that visual cues such as fruit shape and size can strongly influence oviposition behavior in fruit flies (Prokopy et al. 1984, Prokopy & Roitberg 1984). Accordingly, we used 3D-printed fruits to control for shape and size, allowing us to specifically test the effects of independent variation of colour and odour.. These artificial fruits consisted of a black disk of 4 cm in diameter and 1.3 cm in height to hold an odourous liquid, topped by a coloured hemisphere with 32 small perforations for the female to insert her ovipositor. The hemisphere measured 4 cm in diameter and 2 cm in height (Appendix 1). Odourous liquids consisted in 2 mL of 1% water-diluted aroma prepared on the day of the behavioral assays (Appendix 1). Odour release rates were not quantified in the present study. As such, the results should be interpreted in a comparative framework and not directly extrapolated to field conditions. We selected three visual stimuli (blue, yellow, and white) and three natural fruit aromas (cherry, banana, and orange - Culinaide, France) based on literature and preliminary experiments. Although there are some contradictory data (Diaz-Fleischer et al. 1999), we would expect yellow to be the most attractive visual stimuli, white the least attractive and blue intermediate (Katsoyannos et al. 1986). Although “colour” has a specific definition in the context of insect vision (van der Kooi et al. 2021), this term will be used throughout the rest of the manuscript for the sake of simplicity. No study has yet been conducted on natural fruit aromas, but in the field orange is a good, i.e., heavily infested, host. Banana is a bad (rarely infested) host, and cherry is an intermediate host (https://edis.ifas.ufl.edu/publication/IN371, https://www.agric.wa.gov.au/medfly/mediterranean-fruit-fly-host-plants).

### Oviposition behavior assays

The nine colour-odour combinations (Appendix 2), hereafter fruit types, were all tested in four different choice types: no-choice assays and three different one-choice assays. In no-choice assays (homogeneous environment), two identical artificial fruits were placed in each cage. In one-choice assays (heterogeneous environment), the two fruits could differ only in colour (visual choice), only in odour (olfactory choice), or in both colour and odour (visual and olfactory choice). It has to be noted that the two artificial fruits were placed 3 cm apart within the experimental cage. Before being reused, artificial fruits were thoroughly washed with detergent and tap water to remove any residue from *C. capitata* pheromones or fruit aroma.

To estimate oviposition preferences while minimizing both initial behavioural noise and experience-dependent effects, females were tested for the same type of artificial fruits in two standardized 4-hour sessions separated by a two-day interval. The number of eggs laid were summed for each female group over the two testing sessions. Assessing innate oviposition preferences involves a trade-off between short assays, which may be dominated by exploratory or stochastic behaviour, and longer or repeated assays, which may induce learning or physiological constraints such as egg-load depletion. The present design reflects a compromise between these opposing sources of bias. To avoid potential bias related to the placement of artificial fruits in the cages and the positioning of cages within the climate chamber, each fruit type was randomly assigned across all cages, and the position of the fruits within the cage (left or right) was also randomly determined. For the no-choice assays, we conducted 14 replicates for each of the nine fruit types. For the one-choice assays, we performed seven replicates per fruit pair in the visual choice treatment, seven replicates in the odour choice treatment, and eight replicates in the combined visual and odour choice treatment. Because all assays could not be conducted simultaneously, experiments were organized into temporal blocks corresponding to the weeks during which measurements were performed.

The day before the first measurement, females were isolated in experimental cages using a mouth aspirator. In each cage, eight females were placed with water and food (white sugar and inactive dry yeast). On the day of the first measurement, water and food were removed from the cages. The artificial fruits were prepared according to the tested colours and odours and placed in the cages for four hours (from 10 a.m. to 2 p.m). After four hours of oviposition, artificial fruits were removed from the cages and replaced with water and food sources. Then the bases of the artificial fruits containing the laid eggs were photographed and counted automatically with an ad-hoc artificial intelligence model. This model made use of the detectron2 platform (https://github.com/facebookresearch/detectron2) and was trained from a Mask-RCNN ResNet 101 model, using approximately 400 annotated 1000x1000px vignettes (80% used for training and 20% for validation). Annotation consisted of individually masking each egg in the vignette using CVAT v1.1.0. We verified this model on an independent dataset of 49 human-counted photographs, and found to attain satisfying metrics (Kendall’s τ = 0.933; Appendix 3). Two days after the first measurement, the second measurement was conducted following the same protocol.

### Statistical analysis

Statistical analyses were performed using R v.4.3.2 (R core team 2024). All data and R code are available at: https://doi.org/10.57745/8VMCMQ.

Two preliminary checks were first addressed. Firstly, the mortality rate was estimated to check whether experimental manipulations disturbed the individuals. Across all assays, the average female mortality rate between the two measurements of the same block was 1.66%, which is very low, indicating that the females were not particularly stressed during the experiments. Despite low mortality, the number of living females was still accounted as an offset covariate in subsequent statistical analyses. Secondly, a potential left-right bias was also tested for in the no-choice assays. In no-choice assays, there was no significant difference in the numbers of eggs laid in the left and right artificial fruits (paired t-test, t = 0.63, df = 125, p = 0.53), indicating that only colour and odour can affect the number of eggs laid in each of the two fruits per cage.

Then, three response variables were studied: the total number of eggs of the pair of artificial fruits per cage, the number of eggs laid in the focal fruit, and the preference estimated as the number of eggs laid in the focal fruit divided by the total number of eggs of the pair of artificial fruits per cage. Effects of the tested odours and colours were estimated using Generalized Linear Mixed Models (GLMM) with the R package glmmTMB (Brooks et al., 2017), in which blocks were treated as a random effect and the number of females as an offset covariate. The selection of the most appropriate error distribution and link function was based on residual checks provided by the R package DHARMa (Hartig 2024).

To understand the effect of colours and odours on the total number of eggs laid, we first performed a GLMM (family: gaussian) with the fruit colour, odour and the colour-odour interaction in no-choice assays. We then tested for the effect of colours and odours of both artificial fruits on the total number of eggs on the global dataset (no-choice and one-choice assays). Not having any strong *a priori* on the relevant interactions to account for, we applied a systematic model selection procedure on the most complete model (i.e. colour and odour of the focal fruit, colour and odour of the complementary fruit, and all potential interactions) using the package MuMIn (Bartoń, 2024).

Next, to assess the context-dependence of oviposition behavior, the number of eggs laid on the focal fruit was modelled using GLMM (family: quasi-poisson) as resulting from the focal fruit, the choice type and the fruit x choice type interaction. The focal fruit has nine types (3 colours × 3 odours) and the variable “choice type” has four categories (a-no-choice assay, b-visual heterogeneity, c-olfactory heterogeneity, d-visual and olfactory heterogeneity).

Based on the data obtained from one-choice assays, observed preferences were calculated for each pair of artificial fruits using the formula:

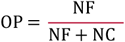

with NF = number of eggs from the focal fruit, and NC = number of eggs from the complementary fruit.

For each pair of fruits presented in one-choice assays, expected preferences, i.e., preferences that should be observed in absence of interaction between fruits, were estimated from the data obtained from no-choice assays. The expected preferences (EP) were calculated using the following formula:

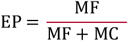

with MF = mean number of eggs from the focal fruit (estimated in no-choice), and MC = mean number of eggs from the complementary fruit (estimated in no-choice).

Preference was modelled as resulting from the effect of the focal fruit, the choice type and their interaction using GLMM (family: ordbeta). We finally analyzed the correlation between expected and observed preferences using a GLMM with observed preference as the response variable, expected preference, choice type and the interaction as explanatory variables using all the heterogeneous assays.

## Results

### Total number of eggs

In no-choice experiments, visual stimuli (χ^2^ = 12.30, df = 2, *p* = 0.002) and olfactory stimuli (χ^2^ = 16.98, df = 2, *p* = 0.0002) significantly influenced the total number of eggs laid. The interaction between visual and olfactory stimuli was not significant (χ^2^ = 2.93, df = 4, *p* = 0.57, Appendix 4). It was thus possible to rank the artificial fruits based on the total number of eggs (Figure 1). Overall, yellow was the colour that most stimulated egg-laying, while white was the least stimulating and blue was intermediate. Cherry and orange odours were not significantly different, and both stimulated egg-laying more than the banana odour (Appendix 5).

**Figure 1.**
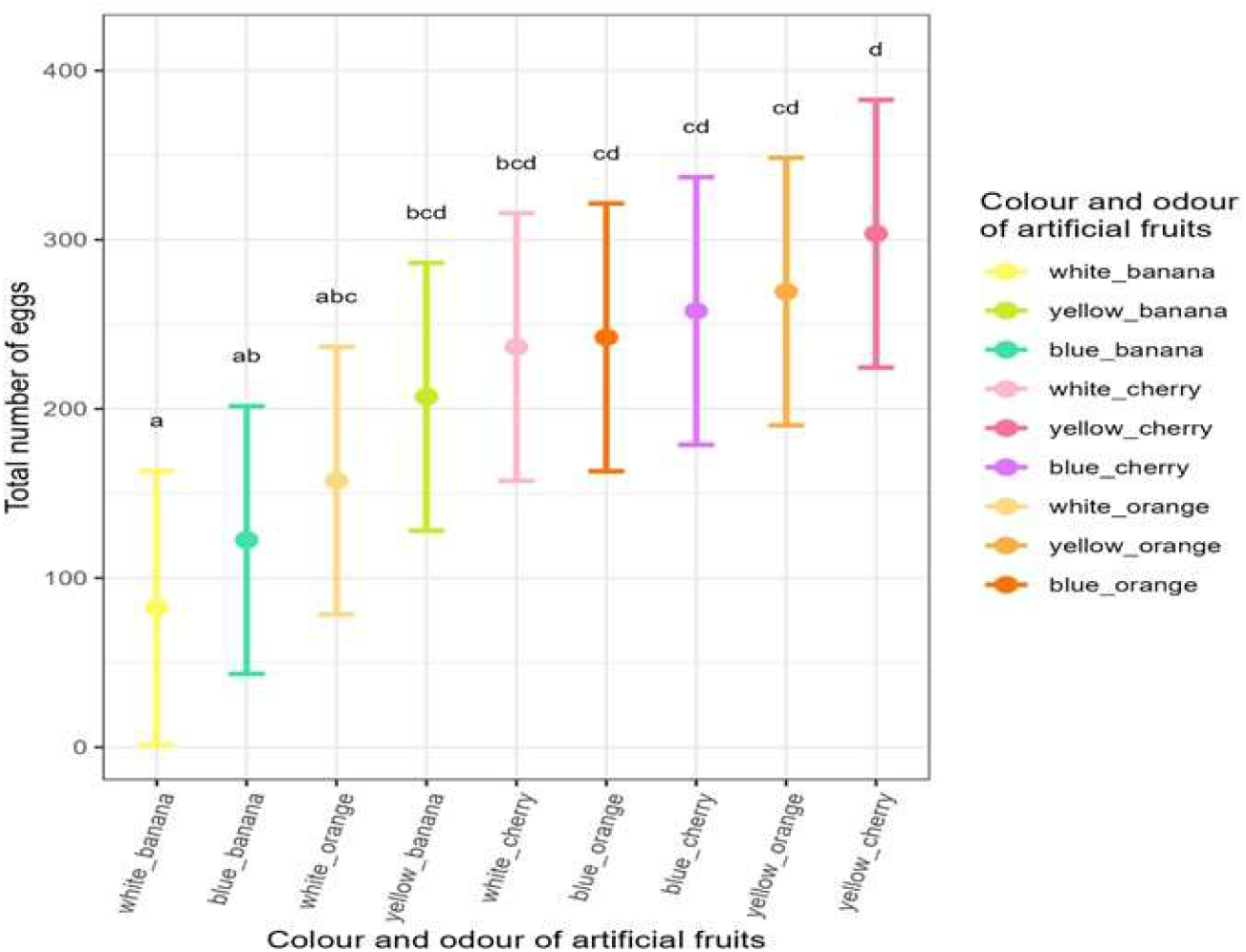
Total number of eggs (with their 95% CI) laid by the eight females during the four-hour assays. The nine colour-odour combinations are ranked in an increasing order. The significance underlying the compact letter display is Bonferroni adjusted.

In the global dataset (no-choice and one-choice assays), the most parsimonious model explaining the total number of eggs laid included a significant effect of the colour and odour of the focal fruit (χ^2^ = 82.09, df = 2, *p* < 2.2.10^-16^ and χ^2^ = 98.81, df = 2, *p* < 2.2.10^-16^, respectively), a significant effect of the colour and odour of the complementary fruit (χ^2^ = 40.82, df = 2, *p* = 1.37.10^-9^ and χ^2^ = 54.97, df = 2, *p* = 1.16.10^-12^, respectively), an interaction between the colour and odour of the focal fruit (χ^2^ = 20.47, df = 4, *p* = 0.0004), an interaction between the colour and odour of the complementary fruit (χ^2^ = 14.68, df = 4, *p* = 0.005), an interaction between the odours of the two fruits (χ^2^ = 80.08, df = 4, *p* < 2.2.10^-16^), and an interaction between the colour of the focal fruit and the odour of the complementary fruit (χ^2^ = 9.57, df = 4, *p* = 0.05, Appendix 6).

### Number of eggs laid in the focal fruit and preference

The number of eggs laid in the focal fruit significantly depended on fruit characteristics (χ^2^ = 21.47, df = 8, *p* = 0.006), on the choice type (χ^2^ = 14.76, df = 3, *p* = 0.002) and their interaction (χ^2^ = 115.15, df = 24, *p* = 7.02.10^-14^, Figure 2, Appendices 7 and 8).

**Figure 2.**
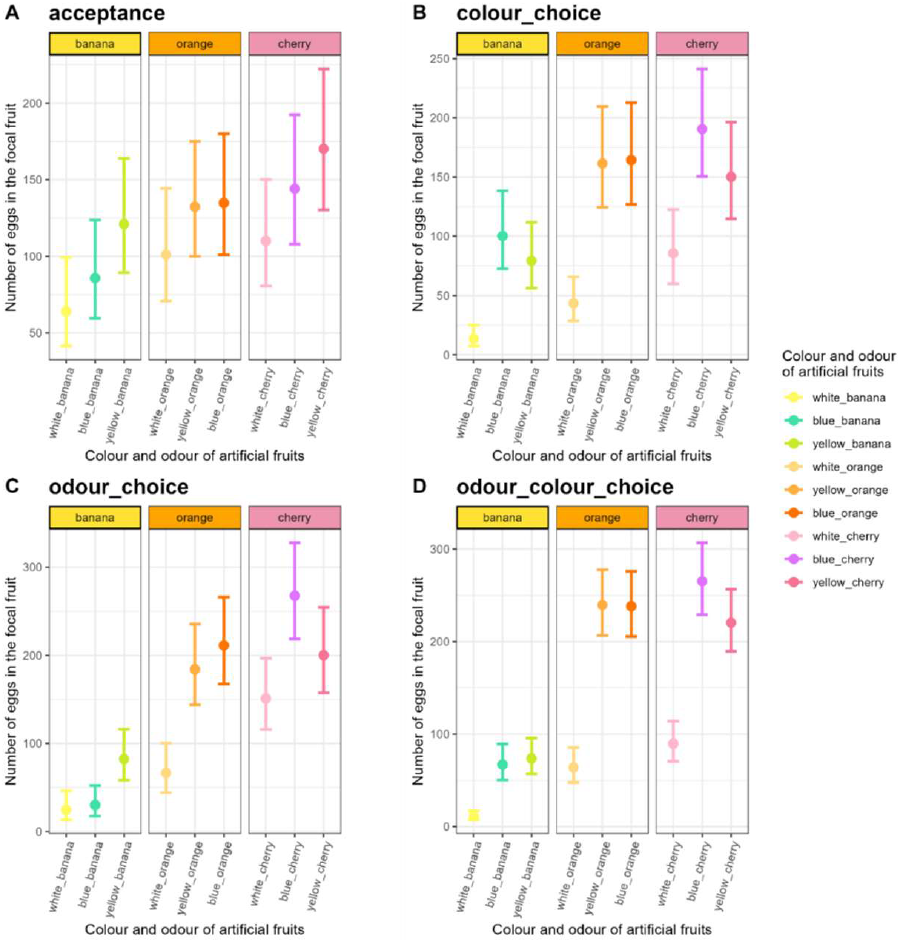
Number of eggs (with their 95% CI) laid by the eight females in the focal during the four-hour assays in the no-choice or acceptance assays (A), in the colour-choice assays (B), in the odour-choice assays (C) and in odour and colour choice assays (D).

The preference was strongly affected by the choice type (χ^2^ = 56.91, df = 3, *p* = 2.7.10^-12^), the interaction between choice type and focal fruit (χ^2^ = 288.30, df = 24, *p* < 2.2.10^-16^) but not by the focal fruit itself (χ^2^ = 3.54, df = 8, *p* = 0.90, Figure 3, Appendix 7).

**Figure 3.**
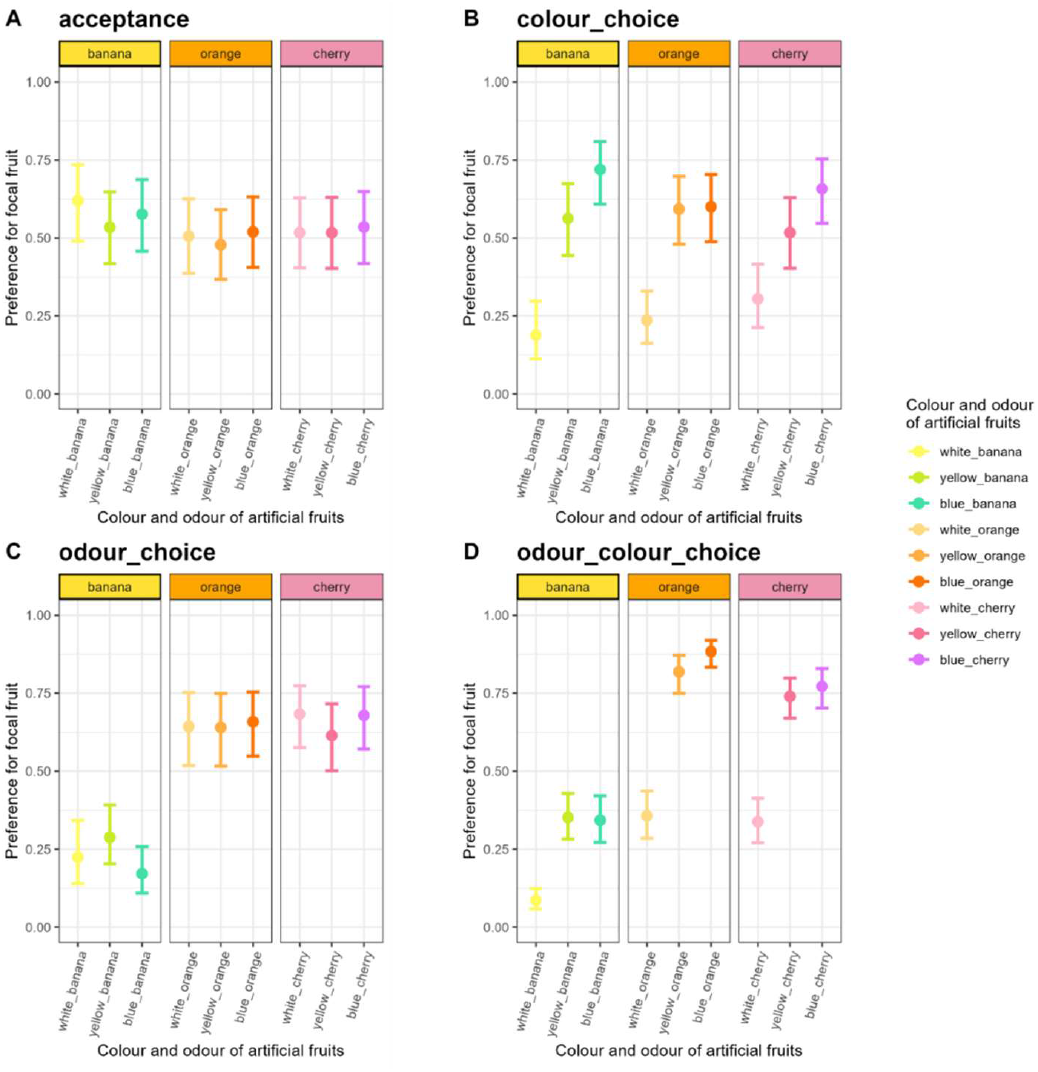
Preference for the focal fruit (with their 95% CI) during the four-hour assays in the no-choice or acceptance assays (A), in the colour-choice assays (B), in the odour-choice assays (C) and in odour and colour choice assays (D).

**Figure 4.**
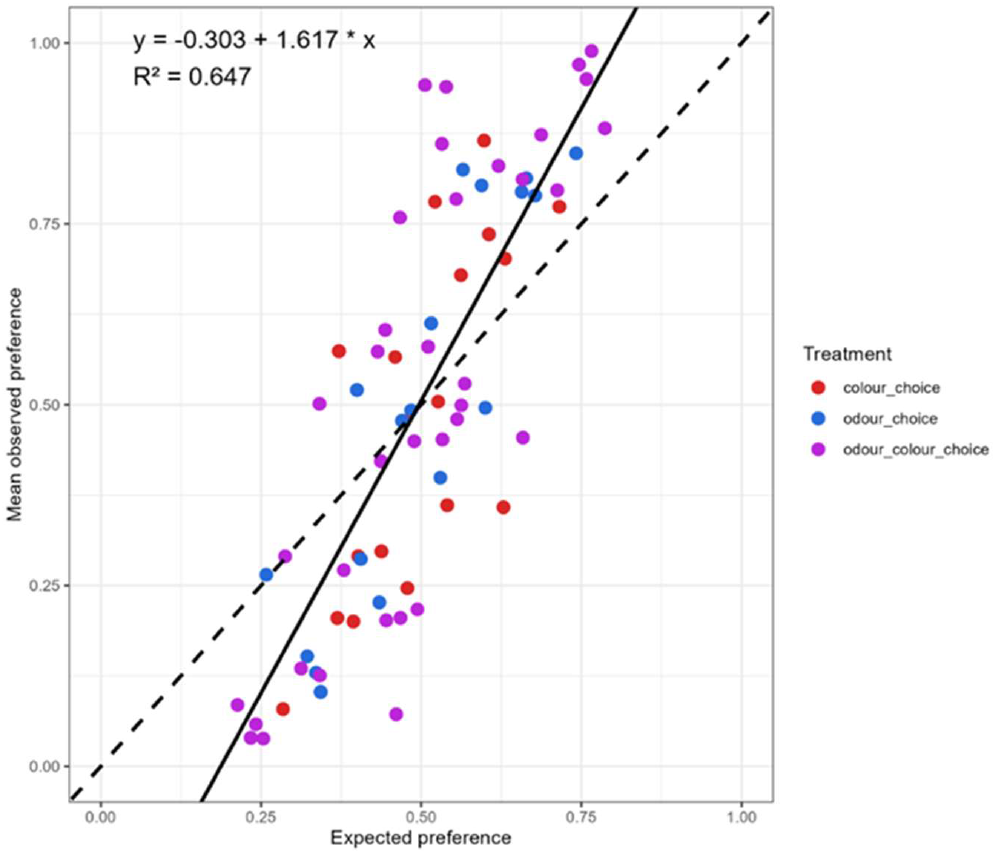
Mean observed preferences versus expected preferences for the three types of choice type. The solid line represents the linear regression fit between observed and expected data and the dotted line represents y=x.

Observed preference was strongly correlated with the expected preference (χ^2^ = 18.49, df = 1, *p* = 1.7.10^-5^), but not by the choice type (χ^2^ = 0.11, df = 2, *p* = 0.95) or the interaction between expected preference and choice type (χ^2^ = 0.13, df = 2, *p* = 0.94, Appendix 9). The observed preferences can be largely explained by the expected preferences, as indicated by the linear regression: *Observed preference* = 1.617 × *expected preference* ™ 0.303 with a high coefficient of determination (R^2^ = 0.647). This relationship implies that when a fruit is preferred in no-choice assays, it is even more preferred in one-choice assays.

## Discussion

Our study adds to the list of works supporting that females of phytophagous insects can alter their oviposition behavior depending on the current availability of different oviposition sites (Jaenike 1978, Schäppers et al. 2017). Here we demonstrate that, in the Mediterranean fruit fly, this alteration stems the high context-dependence of the action of visual and olfactory stimuli on oviposition behavior (Light & Jang, 1996). In no-choice assays, all the nine artificial fruits (three colours × three odours) exceeded the threshold required to stimulate egg-laying, but they displayed sufficiently different visual and olfactory cues to trigger significant different egg-laying outputs (by a factor 1:3 between the two extreme fruits). Our results are in agreement with recent research demonstrating that generalist phytophagous insects have the capacity to rank hosts when given a choice, in contradiction to the “neural limitations” hypothesis postulating that generalist species should not have the neural capacity to identify cues specific to every possible host (Silva & Clarke 2020). In addition to the classification of different colours and odours, it is important to note that the two types of stimuli acted independently of each other in no-choice assays. On the contrary, in the full dataset, including one-choice assays, different types of interactions between these stimuli (visual and olfactory) needed to be accounted for to properly model the data in addition to the simple effect of each stimulus. First, the two types of stimulus (visual and olfactory) significantly interacted within each of the two artificial fruits to explain the number of eggs laid by the female flies. The interaction between the odours of the two fruits and between the colour of the focal fruit and the odour of the complementary fruit also had to be accounted for to best explain the total number of eggs.

A complementary way of analyzing our data is to look specifically at the number of eggs laid in the focal fruit on the overall dataset. As with no-choice assays, the number of eggs laid in the focal fruit in one-choice assays depends on its characteristics in terms of colour and odour, but this time in interaction with the characteristics of the complementary fruit. In other words, the number of eggs laid in a given fruit will be different if the fruit next to it is different from it in odour, colour or both. As expected, we found the same pattern when studying preferences: the effect of stimuli from the focal fruit depends of the female’s evaluation of both stimuli (visual and olfactory) from the complementary fruit. We found that visual and olfactory stimuli do not act independently but interact significantly, influencing oviposition site selection. This finding aligns with studies on sensory integration in insects, particularly flower-visiting species like moths suggesting that while each stimulus (visual or olfactory) can have a direct effect on oviposition behavior, both stimuli are processed together in the brain, contributing to a unified perception of the external environment (Balkenius et al., 2009). This multimodal integration of visual and olfactory stimuli enhances behavioral precision and reflects the adaptive benefits of integrating sensory inputs in a dynamic environment (Hebets & Papaj 2005). The underlying neural mechanisms of this multimodal integration are still not fully understood, but recent studies have shown that responses to combined visual and olfactory stimuli are modulated in the mushroom bodies and at antennal lobe levels of insects (Rennou & Anton 2020). However a study by Vinauger et al. (2019) showed that the modulation of peripheral visual and olfactory sensory stimuli is asymmetric in the mosquito *Aedes aegypti*: CO2 modulates lobula neuropil responses to discrete visual stimuli but visual stimuli do not modulate responses to CO2 in olfactory glomeruli. Our experimental approach allowed us to preserve the integrity of the different sensory channels, which better reflects the natural conditions which insects are confronted to, with multiple simultaneous sensory stimuli. This approach provides a complementary understanding to previous studies that have often isolated sensory stimuli by removing certain modalities (e.g., by ablation of antennae or eye painting; see Riffell, 2020).

Although the female’s response to a given fruit is modulated not only by the fruit individual characteristics (colour and odour) but also by the characteristics of the other available fruit, our study shows that the observed preferences in one-choice are strongly correlated with expected preferences calculated with data from no-choice assays. This result challenges the study by Fanara et al. (2023) that showed a clear decoupling between observed preferences and expected preferences in *Drosophila melanogaster*. Among other existing studies, either the number of fruit pairs tested is too limited to allow statistically robust analyses, or preference is assessed using multi-choice (cafeteria-type) assays. Both approaches limit meaningful quantitative comparisons between acceptance and preference. (Karageorgi et al. 2017, Olazcuaga et al. 2019, Yang et al. 2008, 2015). It would be of great interest to replicate this experimental protocol on a variety of species to better understand the determinants (phylogenetic, ecological, etc.) of the observed patterns. It is also important to note that the relationship between observed preferences and expected preferences is not influenced by the type of heterogeneity between the two fruits (visual, olfactory or both). It is not the nature of the stimulus (visual vs olfactory) that matters, but the relative motivation to lay that the stimulus provides for females after multimodal integration (Renou & Anton 2020). In *C. capitata*, our results show that if the outcome of bimodal interaction between olfactory and visual cues of a fruit is higher than another fruit in no-choice assays, then it will be preferred in one-choice assays by a bias that can be estimated by the coefficients of the linear regression between observed preferences and expected ones. This result suggests that, to be used as a trap plant or in a push-pull, a host plant must elicit more egg-laying than the crop to be protected in no-choice assays. We hope that our results will stimulate new studies that will seek to better understand how the female weigh the various signals to make an appropriate decision for oviposition. Identifying optimal cues in different ecological settings will be key to developing sustainable strategies based on manipulation of insect behaviour.

## Data accessibility

All data and R code are available online at: https://doi.org/10.57745/8VMCMQ

## Supplementary material

Supplementary materials is available online at: https://www.biorxiv.org/content/10.1101/2025.06.20.660717v1

## Acknowledgements

JF and BF acknowledge financial support by the INRAE scientific department SPE and the UMR CBGP from its internal grant support.

