## supplementary material for "Context-dependent oviposition reveals strong association between acceptance and preference in the Mediterranean fruit fly"

**Facon et al.**

**Supplementary material**

Appendix 1:

(A) Artificial fruit produced by 3D printing for studying oviposition behavior of *Ceratitis capitata*. The artificial fruits may vary in colour and odour. (B) Picture of the actual setup for a given pair of artificial fruit (blue vs yellow) inside a 15x15x15cm cage with the actual lighting used during our experiments (light intensity measurement = 4000 ± 200 lux);(C) Reflectance curves of the three types of artificial fruits used in our experiments (white/blue/yellow) in the 350-650nm wavelengths. Darker grey areas delimit the maximal sensitivities of *Ceratitis capitata* (365nm & 485-500nm; Agees 1982)

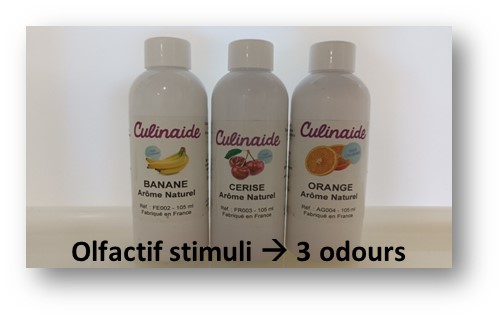

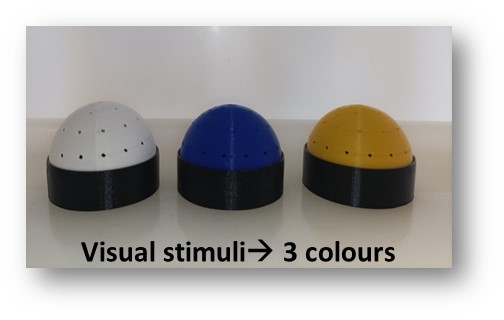
(A)

(B)

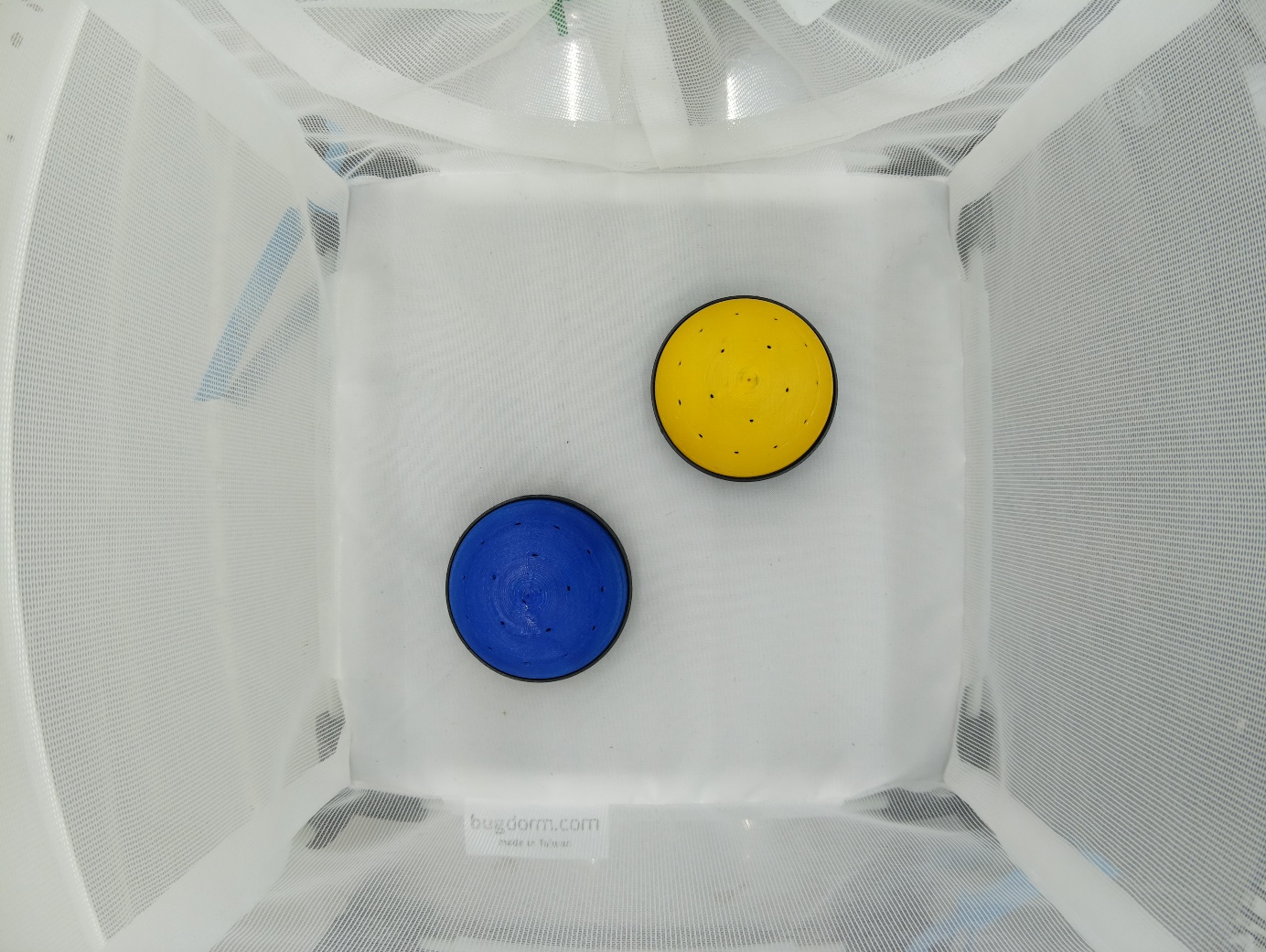

(C)

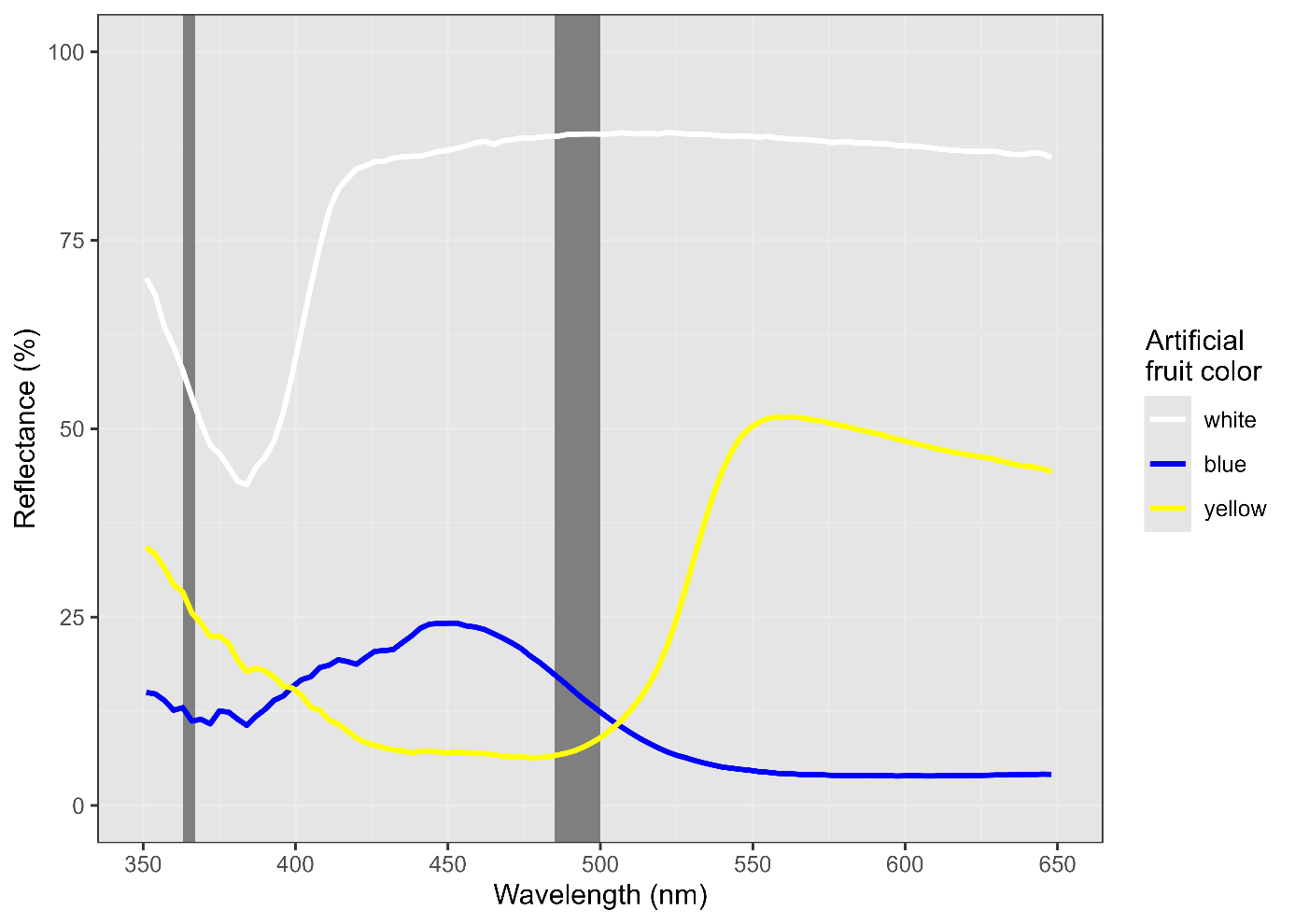

Appendix 2: Numbers of replicates for each combination within the full-factorial design. Red corresponds to no-choice assays with two identical artificial fruits, blue to choice assays between two different colours, orange to choice assays between two different odours and grey to choice assays between two different colours and odours.

|  | 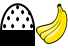 | 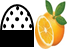 | 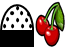 | 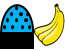 | 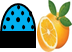 | 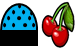 | 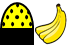 | 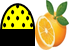 | 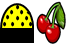 |
| --- | --- | --- | --- | --- | --- | --- | --- | --- | --- |
| 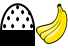 | N = 14 | N = 7 | N = 7 | N = 7 | N= 8 | N= 8 | N = 7 | N= 8 | N= 8 |
| 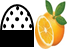 | N = 7 | N = 14 | N = 7 | N= 8 | N = 7 | N= 8 | N= 8 | N = 7 | N= 8 |
| 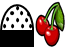 | N = 7 | N = 7 | N = 14 | N= 8 | N= 8 | N = 7 | N= 8 | N= 8 | N = 7 |
| 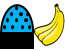 | N = 7 | N= 8 | N= 8 | N = 14 | N = 7 | N = 7 | N = 7 | N= 8 | N= 8 |
| 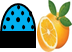 | N= 8 | N = 7 | N= 8 | N = 7 | N = 14 | N = 7 | N= 8 | N = 7 | N= 8 |
| 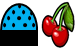 | N= 8 | N= 8 | N = 7 | N = 7 | N = 7 | N = 14 | N= 8 | N= 8 | N = 7 |
| 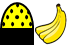 | N = 7 | N= 8 | N= 8 | N = 7 | N= 8 | N= 8 | N = 14 | N = 7 | N = 7 |
| 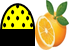 | N= 8 | N = 7 | N= 8 | N= 8 | N = 7 | N= 8 | N = 7 | N = 14 | N = 7 |
| 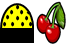 | N= 8 | N= 8 | N = 7 | N= 8 | N= 8 | N = 7 | N = 7 | N = 7 | N = 14 |

Appendix 3: Performance of our custom model for egg-counting. The upper right panel shows the correlation between human and AI counts of eggs for 49 random photographs. Three countings were randomly chosen for their representativeness of low to high counts of eggs (highlighted in red in the upper left panel), and are shown in the remaining A-C panels. In these panels, the prediction of our model is shown through masking of counted eggs (each egg in a different color).

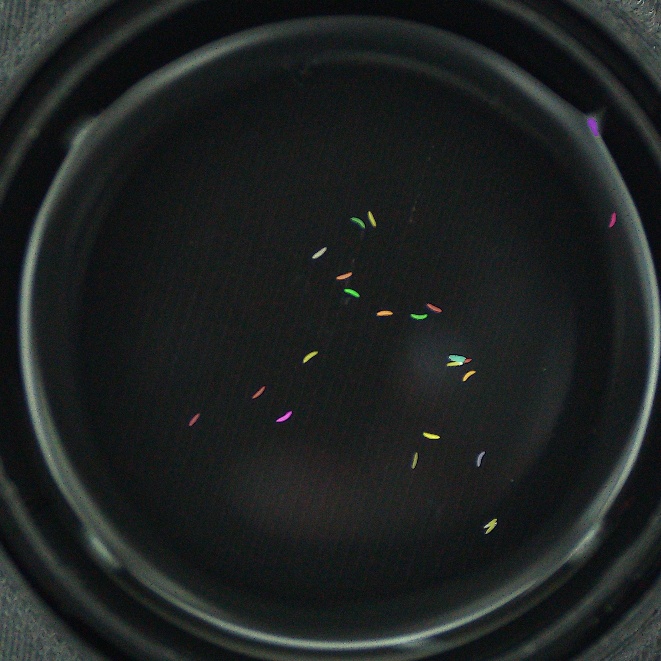

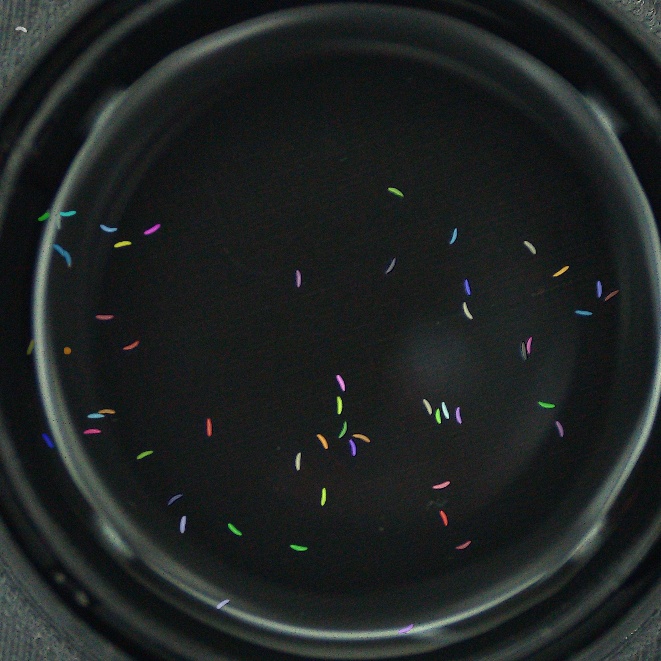

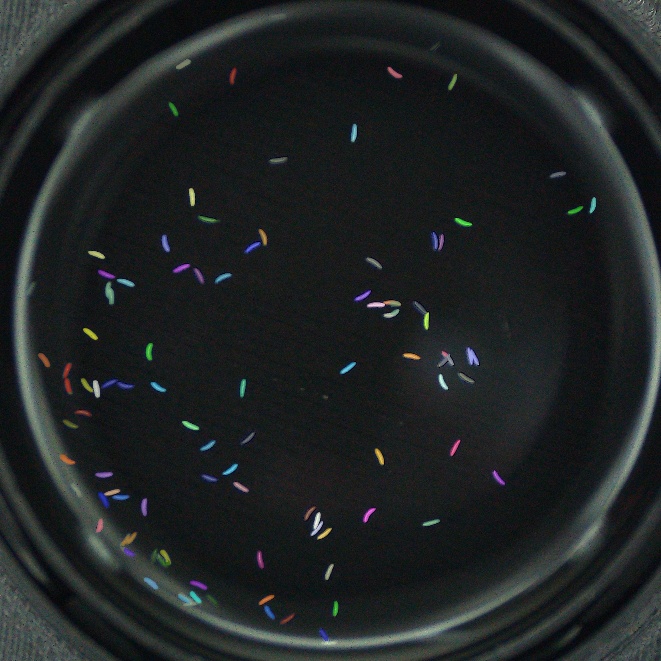

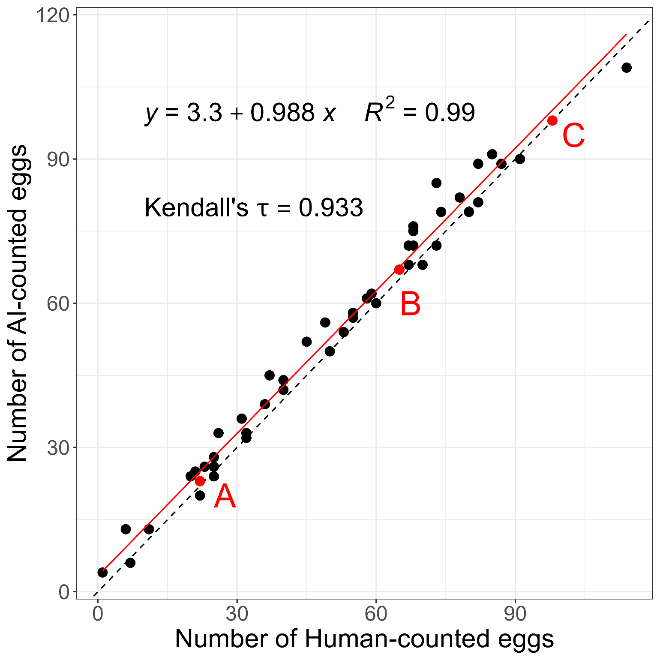

A

C

B

Appendix 4: Results of the GLMM explaining the total number of eggs in no-choice assays by colour of the focal fruit, odour of the focal fruit and their interaction.

| Sources | Df | Chisq | *P*-values |
| --- | --- | --- | --- |
| (Intercept) | 1 | 18.54 | <0.0001 |
| Colour_focal | 2 | 12.30 | 0.002 |
| Odour_focal | 2 | 16.98 | 0.0002 |
| Colour_focal :odour_focal | 4 | 2.93 | 0.57 |

Appendix 5: Pairwise contrast tests on the total number of eggs laid for each pair of colours and each pair of odours.

| Contrast | Estimate | SE | Df | t.ratio | *P*-values |
| --- | --- | --- | --- | --- | --- |
| Colours |  |  |  |  |  |
| Blue-White | 48.7 | 20.9 | 114 | 2.336 | 0.05 |
| Blue-Yellow | -52.5 | 20.8 | 114 | -2.528 | 0.03 |
| White-Yellow | -101.2 | 20.9 | 114 | -4.851 | <0.0001 |
| Odours |  |  |  |  |  |
| Banana-Cherry | -128.7 | 20.9 | 114 | -6.168 | <0.0001 |
| Banana-Orange | -85.8 | 20.9 | 114 | -4.110 | 0.0002 |
| Cherry-Orange | 42.9 | 20.8 | 114 | 2.068 | 0.10 |

Appendix 6: Results of the most parsimonious GLMM (obtained with MuMIn) explaining the total number of eggs laid in the global dataset (no-choice and one-choice assays).

| Sources | Df | Chisq | *P*-values |
| --- | --- | --- | --- |
| Colour_comp | 2 | 40.8170 | <0.0001 |
| Colour_focal | 2 | 82.0875 | <0.0001 |
| Odour_comp | 2 | 54.9688 | <0.0001 |
| Odour_focal | 2 | 98.8064 | <0.0001 |
| Colour_comp:odour_comp | 4 | 14.6763 | 0.005 |
| Colour_focal:odour_comp | 4 | 9.5711 | 0.05 |
| Colour_focal:odour_focal | 4 | 20.4730 | 0.0004 |
| Odour_comp:odour_focal | 4 | 80.0789 | <0.0001 |

Appendix 7: Results of the GLMM explaining the number of eggs laid in the focal fruit (A) and the preference (B) by characteristics of the focal fruit, treatment and their interaction.

| A) Nb_eggs_focal |  |  |  |
| --- | --- | --- | --- |
| Sources | Df | Chisq | *P*-values |
| (Intercept) | 1 | 337.154 | <0.0001 |
| Fruit_focal | 8 | 21.472 | 0.006 |
| Treatment | 3 | 14.755 | 0.002 |
| Fruit_focal :treatment | 24 | 115.150 | <0.0001 |
| B) Preference |  |  |  |
| Sources | Df | Chisq | *P*-values |
| (Intercept) | 1 | 960.4921 | <0.0001 |
| Fruit_focal | 8 | 3.5398 | 0.8961 |
| Treatment | 3 | 56.9130 | <0.0001 |
| Fruit_focal :treatment | 24 | 288.3031 | <0.0001 |

Appendix 8: Scatterplots the number of egg laid for all choices tested during our experiment. Large triangles represent medians, horizontal and vertical segments represent 95% CI,and the black dotted line the x = y line.

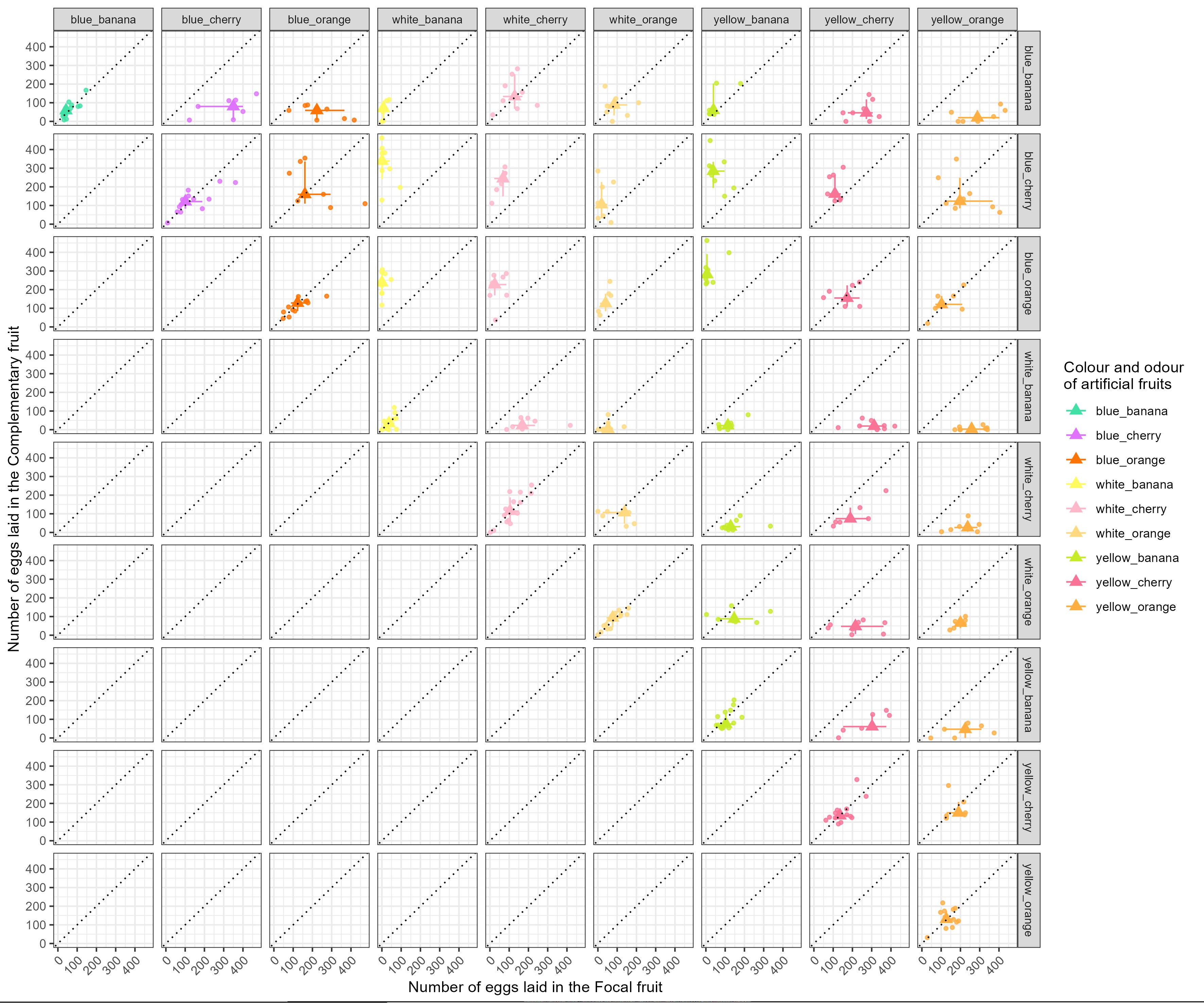

Appendix 9: Results of the GLMM explaining the mean observed preferences by expected preferences, treatment and their interaction.

| Sources | Df | Chisq | *P*-values |
| --- | --- | --- | --- |
| (Intercept) | 1 | 2.3601 | 0.12 |
| Expected_preferences | 1 | 18.4891 | <0.0001 |
| Treatment | 2 | 0.1079 | 0.95 |
| Expected_preferences :treatment | 2 | 0.1292 | 0.94 |
